# Photocrosslinking Activity-Based Probes to Capture the Dynamics of Ubiquitin RING E3 Ligase Interactions

**DOI:** 10.64898/2026.03.13.711684

**Authors:** Sarah F. Chandler, Michael H. Tatham, Emma Branigan, Mark A. Nakasone, Nikolai Makukhin, Alessio Ciulli, Ronald T. Hay

## Abstract

Almost all cellular processes are influenced by ubiquitination. A large family of enzymes known as E3 ligases provide the specificity for ubiquitination, with the largest class among them, the RING E3s, comprising over 600 members in humans. RING E3s facilitate transfer of ubiquitin to substrates by constraining the highly dynamic E2-Ub thioester linkage to be primed for attack from the substrate nucleophile. We have established a workflow that uses a modified ubiquitin carrying a photoactivatable crosslinker that once stably linked to the active site of an E2, creates an activity based probe (ABP) to monitor interactions with E3 ligases. Using this, regions of interaction between ubiquitin and a selection of different RING E3 were determined, which not only confirmed existing structures of E2-Ub-RING-E3 complexes but was also used to assess new Ub-E2-E3 models generated in absence of existing structures.

## INTRODUCTION

Ubiquitination of target proteins requires three classes of enzymes: E1 ubiquitin-activating enzymes, E2 ubiquitin-conjugating enzymes, and E3 ubiquitin ligases^1^. Catalysed by the E1, ubiquitin first forms an ATP-dependent C-terminal adenylate prior to forming a thioester bond to a cysteine in the E1 with release of AMP. A transthiolation reaction delivers the ubiquitin to the catalytic cysteine of a bound E2 to generate a thioester-linked E2-Ub conjugate^2^. This is recruited to an E3 ligase to facilitate the transfer of ubiquitin to substrate^3^. The most common acceptor residue for ubiquitination is lysine, resulting in the formation of an isopeptide bond, although other residues such as serine and threonine^4,5^ can also be modified.

The mechanism of transfer of ubiquitin to substrates from E2-Ub is dependent on the type of E3^6^. One group of ligases known as transthiolating E3s possess a catalytic cysteine, and comprise ∼50 members including HECT (Homologous to the E6-AP Carboxyl Terminus), RBR (RING-between-RING)^7^ and the recently discovered RING-Cys-Relay (RCR) E3s^5^. The largest group of ∼600 consist of the adaptor class of E3s which do not possess a catalytic cysteine and instead catalyse transfer of ubiquitin directly from the E2-Ub to substrate. They are characterised by so-called Really Interesting New Gene (RING) domains which are zinc coordinating scaffolds for the ubiquitin loaded E2^6^. Whilst some RING E3s are active as monomers, most form homodimers or heterodimers^8^. Some RING proteins such as Rbx1 and Rbx2 mediate ubiquitination as components of multi-subunit complexes, such as the cullin-RING ligases (CRL)^8^, which all contain a cullin (Cul1, 2, 3, 4a,4b, 5 or 7) scaffold protein to engage adaptor proteins that bind interchangeable substrate receptors.

RING E3 ligases conformationally constrain E2-Ub so the thioester linkage is primed for attack from the substrate nucleophile, typically the E-amino group of a lysine residue^3^. To achieve this the RING or RINGs contact both ubiquitin and E2, enforcing a ‘closed conformation’^3,9–11^. In absence of E3, the E2-Ub conjugates exist in ‘open conformations’ that are not favourable for transfer to substrate^12^. Understanding the ensemble of Ub-E2-E3 structures by conventional structural techniques is challenging due to their transient nature and conformational flexibility. Photocrosslinking Activity Based Probes (ABP) have promise in this area as they can trap transient conformations^13^. *N*-Maleimido Diazirine (NMD) is a particularly good candidate as it is heterobifunctional and can be linked to a specific site in a ‘probe’ protein through cysteine reaction with the maleimide-thiol and once photoactivated, the carbene group reacts relatively non-specifically with proximal X-H bonds^14^ On this basis we have established a workflow that uses an NMD-linked ubiquitin probe in complex with E2s, that crosslinks from the closed, active conformation to proximal E3 ligases. The identification by mass spectrometry (MS) of cross-linked peptides between ubiquitin and a variety of E3s not only confirmed known structures of E2-Ub-RING complexes but also identified transient complex formations.

## RESULTS

### Design and characterisation of Activity Based Probes for RING E3 ligases

Our E3-targeting ABP design accommodated two important considerations. Firstly, the Ub-E2 was linked via an isopeptide bond rather than the unstable native thioester bond^3^ (Figure 1A). Secondly, the photo-reactive crosslinker should be optimally positioned to capture interacting E3s when the Ub-E2 conjugate is in the closed conformation (Figure 1B). We screened a series of ubiquitin mutants guided by the structure of the RNF4 RING dimer bound to Ub-UbcH5a, to find the optimal cross-linker position. Seven ubiquitin residues at the interface with the RNF4 RING domain (Figure 1C)^3^ were individually mutated to cysteine, expressed and purified (Supplementary Figure 1A-C). Labelling each with NMD produced the expected mass increase of 179 Da (Supplementary Figure 1D), and while some NMD-Ub variants had lower than WT conjugation efficiency in RNF4-dependent 4xSUMO2 conjugation assays^15^ (Supplementary Figure 1E), all were capable of loading onto UbcH5a(C85K) to form NMD-Ub-UbcH5a ABPs (Supplementary Figure 1F). Each of these ABPs was incubated with full-length RNF4 linked to a second RING domain (RNF4+RING), and half of each reaction was UV irradiated, while the other half was not. The products of both were analysed by SDS-PAGE (Figure 1D). While all NMD-Ub variants showed some activity in this assay, D32C appeared the most reactive and was taken forward. A large-scale conjugation and purification of this NMD-Ub(D32C)-UbcH5a(C85K) ABP (herein referred to as NMD-Ub-UbcH5a) was prepared (Figure 2A). Intact MS analysis (Figure 2B + Supplementary Figure 2A-C) showed that while the individual components, NMD-Ub(D32C) and UbcH5a(C85K) were of the expected masses prior to conjugation, the NMD-Ub-UbcH5a product was ∼18 Da larger than expected. Efficient E1-mediated conjugation of ubiquitin to the E2 mutant requires high pH to deprotonate the acceptor lysine. Alkaline conditions can promote hydrolysis of maleimides^16–18^ thus to investigate if this caused the mass anomaly we incubated NMD-labelled Ub(Q31C) at 37°C for 19 hours under conjugation conditions, and analysed the products by intact MS. These conditions generated a +18Da species consistent with maleimide hydrolysis (Supplementary figure 2 D+E). It is thought that hydrolysis of maleimide may stabilize the probe as it opens the ring to form maleamic acid, which prevents the retro-Michael reaction^19^. It is important to account for this mass difference when identifying cross-linked peptides (see below).

**Figure 1.**
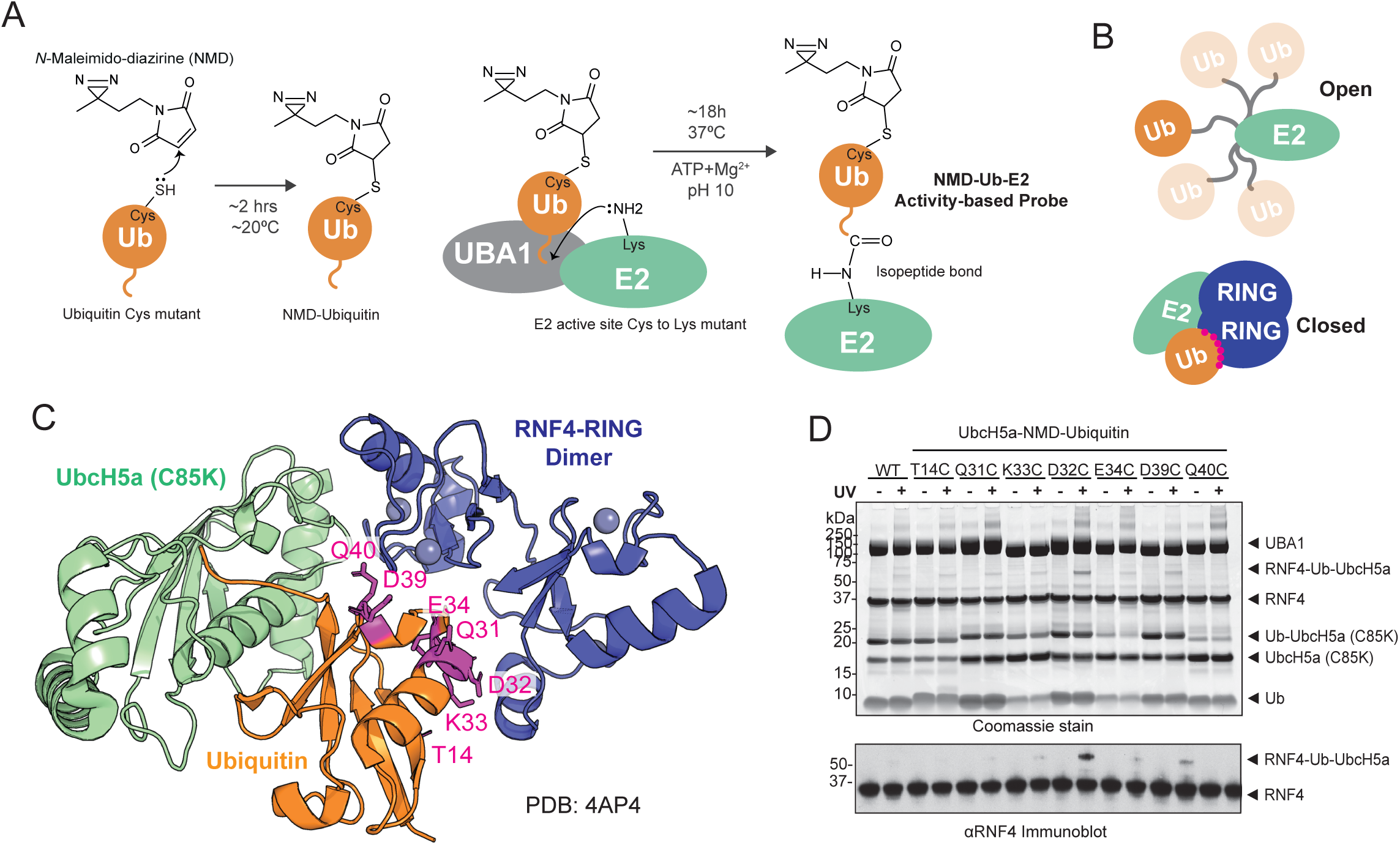
Design of an NMD-labelled Ubiquitin-Directed ABP to capture RING E3s. A. The photoreactive crosslinker N-Maleimido-diazirine (NMD) is linked to a cysteine introduced into ubiquitin by mutagenesis. NMD-Ub is conjugated to the active site of an E2 with the active site cysteine mutated to lysine thereby forming a stable-isopeptide linked conjugate (NMD-Ub-E2) activity-based probe. B. Schematic representation of the open and closed conformations of E2 (green)-Ubiquitin (orange) with E3 RINGs (dark blue). C. Crystal structure of RNF4 RING dimer (blue) bound to ubiquitin (orange) conjugated to UbcH5a C85K mutant (pale green) in the closed conformation (PDB:4AP4). Seven sites for photocrosslinker coupling are indicated (magenta). D. Photocrosslinking of the seven different ubiquitin mutants in NMD-Ub-E2 ABP with RNF4+RING. Proteins were separated by SDS-PAGE and visualised with Coomassie blue staining (top) or immunoblotted for RNF4 (bottom).

**Figure 2.**
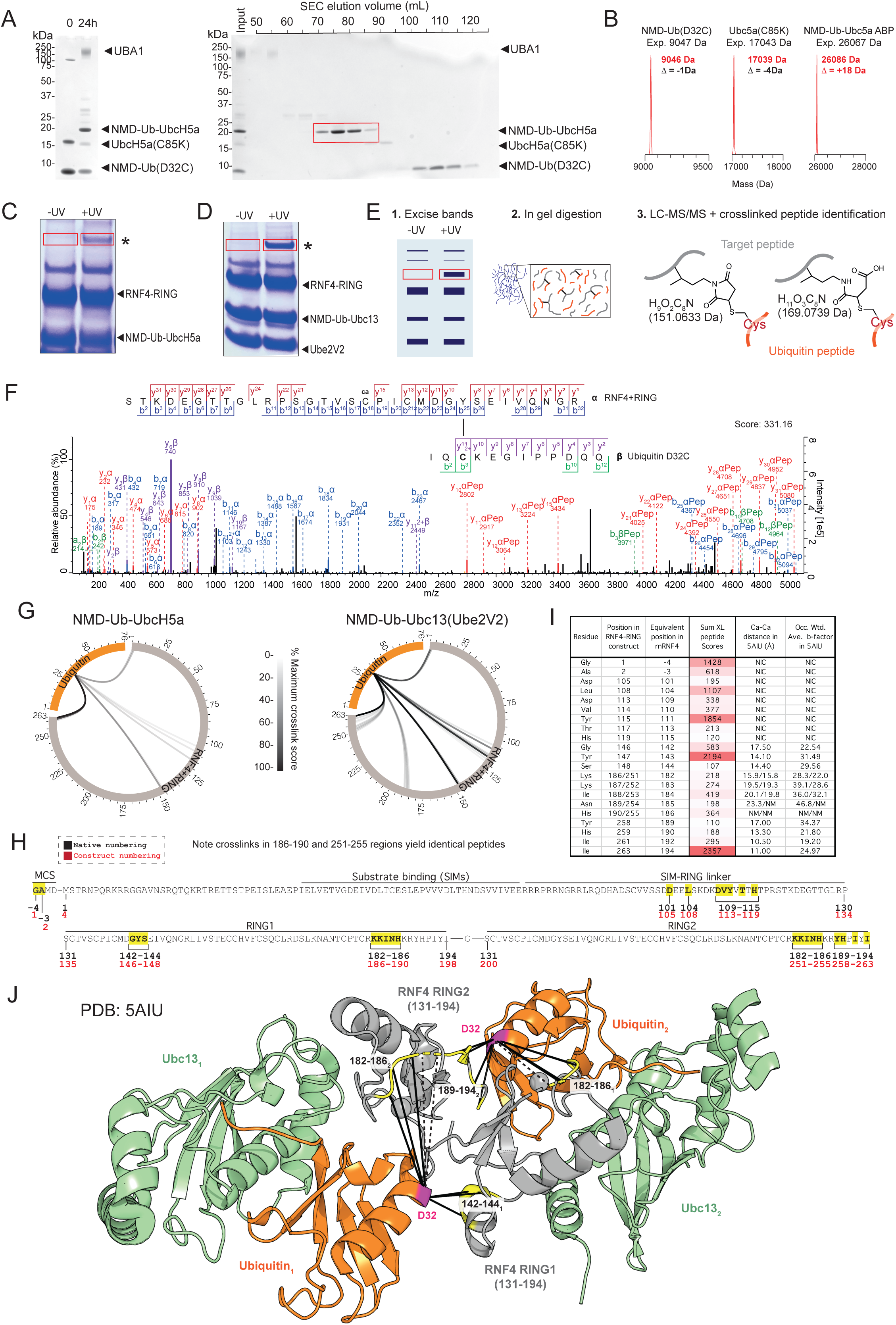
NMD-Ubiquitin-E2 based photoreactive probes confirm the in-solution structure of RNF4+RING bound by ubiquitin loaded E2. A.(Left) SDS-PAGE fractionation of conjugation reactions containing NMD-ubiquitin (D32C), UbcH5a (C85K) and UBA1. Reaction products were separated by size exclusion chromatography (right) and fractions containing purified NMD-Ub-UbcH5a were pooled (red). B. LC-MS analysis of NMD-ubiquitin (D32C), UbcH5a (C85K) and NMD-Ub-UbcH5a. C+D. SDS-PAGE analysis of the products of photocrosslinking (+UV) or control reactions (-UV) containing RNF4+RING with either NMD-Ub-UbcH5a (C) or NMD-Ub-Ubc13 (D). E. Workflow for identification of sites of crosslinking by mass spectrometry. F. Exemplar MS/MS spectrum for crosslinked peptide linking RNF4 Y143 to ubiquitin D32C. α peptide from RNF4 and β peptide from ubiquitin D32C G. Circular XiView plots showing crosslinks between RNF4+RING (grey) and ubiquitin (orange) using ABPs containing UbcH5a (left) or Ubc13 (right). Each line represents a unique crosslinked pair of residues with lines shaded by cumulative score relative to the highest (100% black). H) Crosslinked amino-acids (yellow) from the RNF4+RING construct comprising full-length RNF4 fused to a second RING domain. Corresponding distances between alpha carbons (I) based on the published structure of Ub-Ubc13 bound to an RNF4 RING dimer (PDB: 5AIU). Sum of the scores of all crosslinked peptides that evidenced links between UbC32 and the specified E3 residue are shown along with occupancy weighted b-factors from the structure. (J). Identified cross-links (solid lines) shown on the Ub-Ubc13-RNF4+RING structure (5AIU). Crosslinked regions shown in yellow. Broken lines show cross-links to unmodeled regions.

### Photocrosslinking with RNF4 confirms existing structures and identifies transient contacts between ubiquitin and non-RING domains

To broaden the scope of this study, we generated a second probe, NMD-Ub-Ubc13 ABP by conjugating NMD-labelled ubiquitin (D32C) to Ubc13 (C87K/K92A) (Supplementary Figure 3A). The two ABPs, NMD-Ub-UbcH5a and NMD-Ub-Ubc13 (combined with its heterodimeric partner, Ube2V2) were tested for crosslinking to the RNF4+RING construct. Both formed UV-dependent higher molecular weight adducts (Figure 2C, D) which were excised from the gel along with equivalent sections from the control lanes and tryptic peptides analysed by LC-MS/MS (Figure 2E). Protein intensity data confirmed that UV exposure increased amounts of E2s, RNF4 and ubiquitin in the +UV samples (Supplementary Figure 3B, C), consistent with the formation of Ub-E2-RNF4 covalent crosslinks. Crosslinked peptides are often large with complex MS/MS spectra. Data analysis can be hindered by the combinatorial explosion in search space caused by considering all possible peptide pairs during PSM matching^20^ . This is reduced as the crosslinkers were restricted at one end to cysteine reactivity (maleimide) but were unrestricted at the other (diazirine). The data were searched for both non-hydrolysed NMD (151.0633 Da) and the hydrolysed form (169.0739 Da) (Figure 2E) but only one crosslink involving the non-hydrolysed form remained after data filtering, and was the only NMD-151 crosslink identified between Ub and an E3 among all the experiments described in this study (see Supplementary datafile 1). The remainder of this report will focus on NMD-169 crosslinks. Figure 2F shows an example of a high-scoring cross-linked peptide. Broad agreement was apparent for both probes (Figure 2G). Ubc13 provided the largest dataset of 21 crosslinks, with many being within the two RINGs (Figure 2H). Mapping these onto the published structure of Ub-UBC13 bound to two RNF4 RINGs (PDB: 5AIU) showed most crosslinked residues were 10-20Å away from ubiquitin D32 although His186 was not modelled in this structure (Figure 2 I, J). While all of these distances are beyond the nominal length of the NMD linker (8-10Å), these Cα-Cα measurements do not account for sidechain lengths or the precise point of reaction of the diazirine on RNF4. Even so, some crosslinked residues including Lys183, Ile184 and Asn185 appeared too far away from UbD32 (19.5, 20.1 and 23.3Å respectively). However, the b-factors of these residues in this structure are relatively high (Figure 2I), suggesting these regions of RNF4 are relatively flexible and in some conformations may be closer to UbD32 than shown in this structure.

Some high-scoring crosslinks were also identified outside the RING domains. Crosslinks between the region linking the substrate-binding and RING domains indicate that these regions contact the E2∼Ub. This is supported by single molecule FRET experiments suggesting that the linker region, although disordered (Supplementary Figure 3D) comes into close proximity to the RING (Murphy et al., 2020) where it could bring the bound substrate towards the E2∼Ub to facilitate substrate modification. The data for the NMD-Ub-UbcH5a probe was also consistent with the published structure in the resolved regions (Supplementary figure 3E, F, G) albeit with fewer identified links.

### Mapping Ubiquitin interactions with the heterodimeric RING E3 RNF2-BMI1

Having optimised our ABPs with the RNF4 double RING construct we then sought to probe a heterodimeric E3 ligase. RNF2-BMI1 bound to UbcH5c has been structurally resolved^21^, although ubiquitin was not included in the complex. We therefore undertook a crosslinking analysis using NMD-Ub-UbcH5a to interrogate a model generated by docking ubiquitin into this structure. UV exposed and control reactions were prepared (Supplementary Figure 4A), and protein intensity data showed crosslinking had occurred (Supplementary Figure 4B). Six unique crosslinks were identified (Figure 3A-C), of which 5 were to BMI1, and one in RNF2. Notably, both RNF2 and BMI1 formed crosslinks via an N-terminal serine derived from the expression constructs (Figure 3B), neither of which were present in the published crystal structure (Figure 3C). According to the model generated by docking ubiquitin into the closed conformation in the existing structure (Figure 3D) the remaining crosslinks to resides Lys73, Leu75 and Asp77 in BMI1 were all <18Å in distance from ubiquitin D32 (Figure 3C, D), supporting this as a good representation of the complex in solution. An Alphafold model based on the RNF2, BMI1 ubiquitin and UbcH5a constructs used in our crosslinking experiment, compared favourably to the docked ubiquitin structure (Supplementary Figure 4C), and confirmed the N-terminal regions of RNF2 and BMI1 are flexible, thus explaining their cross-linking with ubiquitin(C32). Gly109 in BMI1 was also missing from the crystal structure but simple extension of the pre-existing α-helix would take it well beyond the range of the crosslinker. However, the AlphaFold model suggests that Gly109 sits at the end of a helical turn (Supplementary Figure 4C) only 12.2A from UbC32, although Alphafold pLDDT scores in this region are relatively low.

**Figure 3.**
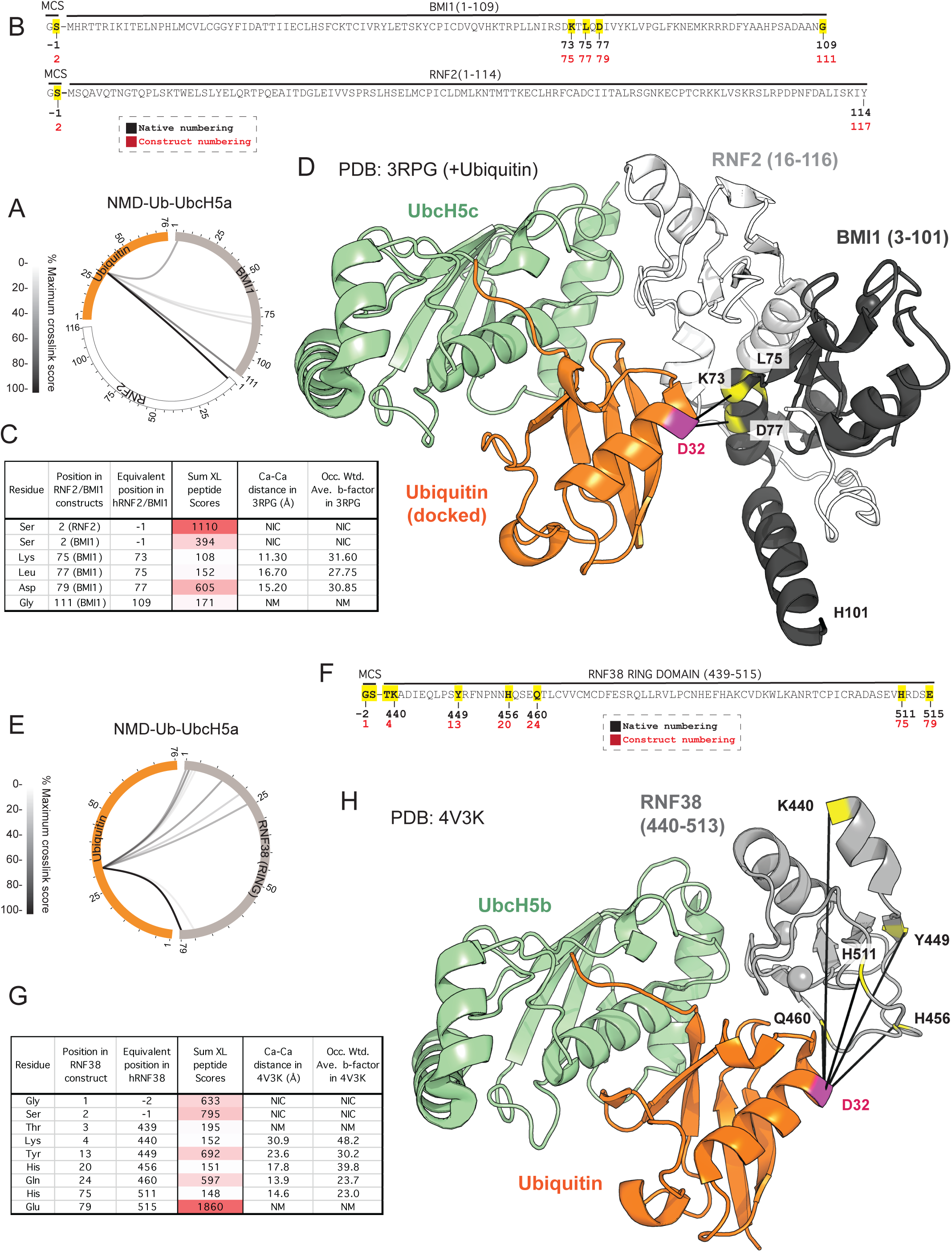
NMD-Ub-UbcH5a ABP captures the heterodimeric RING E3 RNF2/BMI1 and the monomeric RING E3 RNF38. A. Circular XiView plot showing crosslinks between BMI1 (grey) or RNF2 (white) and ubiquitin (orange) using the NMD-Ub-UbcH5a ABP. Each line represents a unique crosslinked pair of residues with the line density showing the relative sum of the scores for each. Sequences and sites of crosslinks (yellow) are shown in B. C. Corresponding distances between alpha carbons based on the model in D. Sum of the scores of all crosslinked peptides that evidenced links between UbC32 and the specified E3 residue are shown along with occupancy weighted b-factors from the structure. D. Published structure of UbcH5a-BMI1-RNF2 (PDB: 3RPG) with ubiquitin docked in the closed conformation. E. Circular XiView plot showing crosslinks between RNF38 (grey) and ubiquitin (orange) using the NMD-Ub-UbcH5a ABP. F. Amino-acid sequence of the RNF38 (439-515) construct with cross-linked residues (yellow). Distances between alpha carbons calculated (G) based on the Ub-UbcH5b-RNF38 structure (PDB: 4V3K). Sum of the scores of all crosslinked peptides that evidenced links between UbC32 and the specified E3 residue are shown along with occupancy weighted b-factors from the structure. H. PDB: 4V3K with identified crosslinks shown.

### RNF38 photocrosslinking with NMD-Ub-UbcH5a confirms the existing structure and identifies mobile regions of the ligase

We next tested the monomeric RING ligase RNF38 which has been resolved by crystallography in complex with Ub-UbcH5b^22^. We performed a photo-crosslinking assay using the NMD-Ub-UbcH5a probe with the RING domain of RNF38 (439-515) (Supplementary Figure 4D, E) and found 9 high-scoring crosslinks (Figure 3E-G). Again, crosslinks to N- and C-terminal amino-acids were apparent, although they were not present in the published structure (Figure 3H). Of those that were resolved, H456, Q460, and H511 were <18Å from D32 of ubiquitin. K440 and Y449 were 30.9 and 23.6Å away, although, as was the case for RNF4, the b-factors of these residues implies structural flexibility (Supplementary Figure 4F) and are likely to be relatively mobile.

### Crosslinking evidence that Ub-UbcH5a can bind a symmetrical CHIP dimer

Our ABPs provided crosslink data with various RING containing E3s. To determine if we could employ the ABP to analyse interactions with non-RING E3s we tested the U-box E3 ligase CHIP. U-boxes have similar folds to RING domains but lack zinc ions and are instead stabilized by hydrogen bonds and salt-bridges^7^ . Full-length human CHIP was crosslinked to the NMD-Ub-UbcH5a probe (Supplementary Figure 5 A, B), resulting in identification of 8 crosslinks (Figure 4A, B). Consistent with the other E3s tested, the N-terminus of CHIP formed a crosslink to UbC32. Five crosslinks mapped to the U-box domain (E242, P243, E258, E259 and H266), one to the TPR domain (Y121), and one to the protomer crossover helix (S191). Although no human CHIP-E2-Ub complex structure is available, a crystal structure of the apo mouse CHIP shows the E3 forms an asymmetrical homodimer composed of an ‘elongated’ protomer with a long linear helical domain projecting away from the U-box domain, and a ‘compact’ protomer that has a helical domain with a break half way along its length causing a bend^23^. This break twists the TPR and U-box domains out of alignment, meaning only one Ub-E2 thioester can bind to the ‘elongated’ protomer. To assess our crosslinking data, we recapitulated a Uev1a-Ub-Ubc13-mCHIP dimer by combining the two published structures of Uev1a-Ubc13+mCHIP(U-Box domain) (PDB: 2C2V) with the full-length CHIP dimer (PDB: 2C2L) and docked ubiquitin into the closed conformation on Ubc13 (Figure 4D). In this composite structure most of the interaction between the Ub-Ubc13 occurs with the U-box domain of the ‘elongated’ CHIP protomer. The crosslinks identified were closer to the U-box of the other ‘compact’ CHIP protomer (Figure 4D). Crosslinks to E243 and P244 in the U-box α-helix are 12.6 and 13.6Å respectively from UbD32 (Figure 4C, D). However, crosslinks to E259 and E260, also in a helical region, are both over 20Å away from ubiquitin D32 (Figure 4C, D). Furthermore H267 (predicted to be in a disordered U-box region) and Y122 in a TPR helix are both ∼30Å away from UbD32. Due to relatively low resolution of the mCHIP dimer structure, flexibility could not be assessed by b-factor so an alphafold model was generated using two copies of mCHIP (24-304) and one each of Uev1a, Ubc13 and ubiquitin, and the pLDDT used as a proxy for disorder propensity (Figure 4E). While the positioning of Ub-Ubc13 and one CHIP protomer was consistent with the composite structure, Alphafold predicted a symmetrical rather than an asymmetric CHIP dimer (Figures 4D, E). In this symmetrical dimer the U-box and TPR domains are considerably closer to ubiquitin D32 than in the crystal composite model, and all predicted crosslinker distances are shorter (Figure 4C). The most striking difference is Y122 which is over 18Å closer to ubiquitin D32 in the symmetrical model. These data suggest that a symmetrical CHIP dimer may exist in solution, which could bind two copies of ubiquitin-loaded E2s.

**Figure 4.**
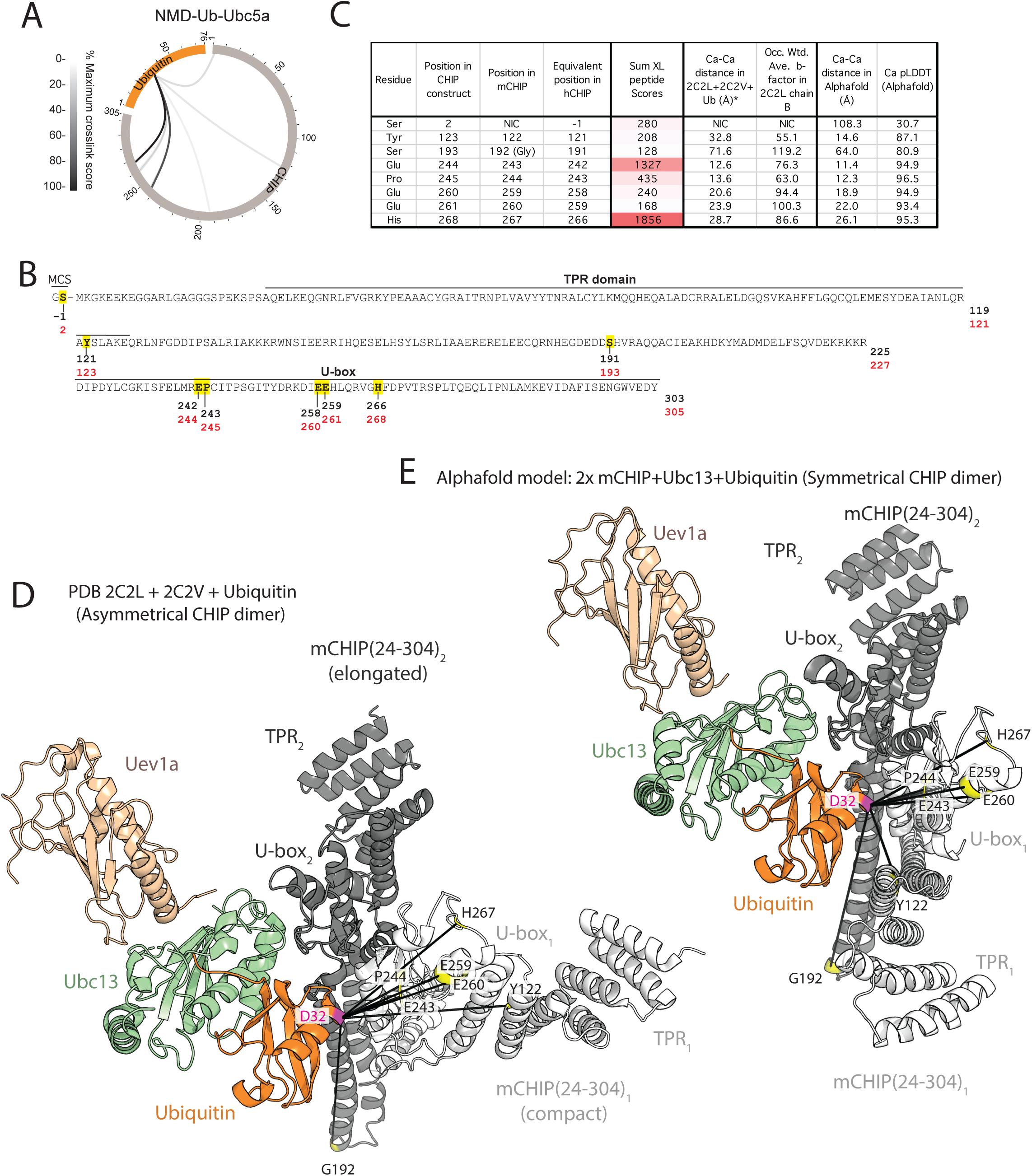
Crosslinking between NMD-Ub-UbcH5a and CHIP support a symmetrical dimer binding arrangement. *A.* XiView summary for crosslinks between NMD-Ub-UbcH5a and hCHIP. B. Sequence of CHIP used in this study with yellow residues showing sites of crosslinking. C. Distances measured between alpha carbons of ubiquitin C32 and cross-linked residues from CHIP according to the models shown in D and E. Sum of the scores of all crosslinked peptides that evidenced links between UbC32 and the specified E3 residue are shown along with occupancy weighted b-factors from the structure or pLDDT scores from Alphafold. (D) Model generated by docking Ubc13/Uev1a from PDB: 2C2V onto the crystal structure of the apo full-length mouse CHIP asymmetrical dimer PDB: 2C2L by aligning U-box domains, with a single ubiquitin (D32C) is docked in the closed conformation. E. Alphafold model of two copies of mouse CHIP with one each of Ubc13 and ubiquitin.

A second Alphafold model was generated using two copies of each of the proteins used in the crosslinking study; hCHIP(1-303), UbcH5a and Ubiquitin (Supplementary Figure 5C-E). This confirmed that the N-terminal crosslink is likely to be due to high mobility and intrinsic disorder of the CHIP N-terminal domain (Supplementary 5C). It also showed that the TPR and U-box domains are confidently modelled and support a 2:2 (CHIP:Ub-E2) stoichiometry complex (Supplementary Figure 5D). A recent cryoEM study using an FAb that interacts with the E2 binding site in the CHIP U-box^24^ found CHIP can adopt a symmetrical structure. The symmetrical structure is also supported by MD simulations^25^ and the structure of a TPR depleted version of CHIP ^26^.

## DISCUSSION

Understanding transient conformations of protein complexes poses a challenge to traditional structural techniques. Activity based probes have great potential in this endeavour as they can provide information on the broader ensemble of protein complex formations in solution. While some ABPs have been reported for ubiquitin RING E3s^27^ and SUMO E3s^28^, they have not been used to provide detailed structural mapping. We have developed a photocrosslinking ABP based on an E2 enzyme linked via an isopeptide bond to ubiquitin carrying an NMD crosslinker and used it to precisely map sites of interaction with multiple RING E3 types. In all experiments, cross-links were identified that were consistent with the known crystal structures or complexes modelled from crystal structures.

Many well-evidenced cross-links were not apparently consistent with published structures. Notably, in all experiments, residues close to the N-termini of the E3 constructs formed cross-links with the probe, highlighting flexible or disordered protein termini as common crosslinking sites by virtue of ensemble proximity^29,30^, rather than fixed proximity. Even within the structured RING domains of the E3s, cross-links were found that, according to the published crystal structures, were too distant to be linked via NMD. However, in most cases these were localised to regions of elevated b-factor (Supplementary Figure 5F), and therefore, these apparent distance violations were likely evidence of flexibility and indicate regions of the E3 that adopt multiple conformations in solution.

A potential application of this technology is the assessment of protein complexes modelled in silico. The most widely used tool for structural modelling, Alphafold, reports pLDDT (Predicted Local Distance Difference Test) values for all residues in a model, which represents confidence in the residue position shown in a model. We generated multiple Ub-E2-E3 Alphafold models based on the proteins used in the cross-linking experiments (Supplementary Figures 3D, 4C and 5C) and observed a strong inverse correlation between Cα-Cα distance and pLDDT (Supplementary Figure 5G). Therefore, similarly to the crystal structure b-factors (Supplementary Figure 5F), longer Cα-Cα distances are indicative of flexible or disordered domains that may adopt conformations not presented in the models. Based on these observations we advocate the combined use of crosslinking experiments with in silico modelling to better understand the dynamics of Ub-E2-E3 complexes. For the NMD crosslinker, our data suggest Cα-Cα distances below 40Å in combination with Alphafold pLDDT scores greater than 70 (supplementary Figure 5G) are indicative of residues close to the probe in most conformations within the ensemble. Longer crosslinks with lower pLDDT scores indicate residues that are only close to the ABP in a minority of the total ensemble of conformations (Supplementary Figure 5G). Crosslinks violating this relationship may also have some bearing on the complex dynamics. For example, S191 in hCHIP was found cross-linked to UbC32 over 70 Å away in the static model (Supplementary Figure 5E), but with a relatively high pLDDT score of 81. S191 is at the very apex of the helical crossover region of CHIP and is significantly different between the elongated and compact forms. However, even in the compact conformation S191 is still >60Å from UbC32, suggesting that this crosslink forms when CHIP is in a conformation not shown by either the symmetrical or asymmetrical dimer, perhaps during the process of flipping between the two, when MD simulations suggest that the helix could unwind^25^.

In summary, this work demonstrates the successful application of NMD-ABPs to exploring they dynamics Ub-E3-E3 complexes. In testing the utility of our method, we focused on a single ubiquitin mutant (D32C) that was most effective at crosslinking to RNF4. It is possible that crosslinking studies not involving RING E3s may require screening of ubiquitin mutants to determine the best NMD positioning. We also expect that NMD-based crosslinking studies combined with the workflow described here are likely to have utility extending to all ubiquitin-like proteins.

## Materials and Methods

Expression and purification of RNF4+RING, Ubiquitin, UbcH5a(C85K), Ubc13(C87K/K92A) and Ube2V2.

Proteins were expressed in *Escherichia coli* BL21(DE3). Starter cultures (10–25 mL) of LB medium supplemented with appropriate antibiotic were inoculated from freshly streaked LB agar plates and incubated overnight at 37 °C with shaking at 220 rpm. 5 mL overnight culture was used to inoculate 500 mL of LB medium containing appropriate antibiotic in a 2 L flask. Cultures were grown at 37 °C with shaking at 220 rpm until an OD600 ∼0.6–0.8 was reached. Cultures were cooled on ice for 10–20 min prior to induction with 100 μM IPTG at 20 °C with shaking at 220 rpm for 17–19 h. Bacterial cell pellets were resuspended in lysis buffer consisting of 50 mM Tris, 250 mM NaCl, and 10 mM imidazole, adjusted to pH 7.5, and supplemented with EDTA-free complete protease inhibitor cocktail (Roche), using 40 mL buffer per 0.5 L of cell culture. Cell suspensions were lysed by sonication (Digital Sonifier, Branson), and insoluble material was removed by centrifugation (27,200 g, 45 min, 4 °C). The clarified lysate was filtered through a 0.2 μm filter and loaded onto a Ni–NTA agarose column (Qiagen) using 1 mL resin per 0.5 L culture, pre-equilibrated with binding buffer containing 50 mM Tris, 250 mM NaCl, and 10 mM imidazole, pH 7.5. The column was washed sequentially with approximately 8 column volumes of binding buffer followed by 6–8 column volumes of washing buffer containing 50 mM Tris, 250 mM NaCl, and 30 mM imidazole, pH 7.5. Bound proteins were eluted using 50 mM Tris, 150 mM NaCl, 150 mM imidazole, and 0.5 mM TCEP, pH 7.5. Pooled Ni–NTA elution fractions were dialyzed overnight at 4 °C against buffer containing 50 mM Tris, 150 mM NaCl, and 0.5 mM TCEP after addition of 1 mg TEV protease per 10 mg of 6His-tagged ubiquitin or 1 mg TEV protease per 50 mg for all other 6His-tagged proteins. For incomplete cleavages, additional TEV protease was added and samples were incubated at room temperature for approximately 4 h. After cleavage, imidazole was added to a final concentration of 10 mM and samples were centrifuged (3,900 g, 15 min, 4 °C) to remove precipitated material. The clarified supernatant was applied to a Ni–NTA agarose column pre-equilibrated with buffer containing 50 mM Tris, 150 mM NaCl, 10 mM imidazole, and 0.5 mM TCEP, pH 7.5. The flow-through fraction was collected and if homogeneous, samples were dialyzed against 50 mM Tris, 150 mM NaCl, and 0.5 mM TCEP, pH 7.5, and concentrated to approximately 500 μM using Vivaspin centrifugal concentrators (Sartorius; 3–10 kDa MWCO) at 3,000 g and 4 °C, and aliquots stored at −80 °C. Where necessary further purification by size-exclusion chromatography was performed. SDS-PAGE followed by Coomassie blue staining was used to assess fractions and those containing homogenous protein were pooled, concentrated, flash-frozen in liquid nitrogen and stored at −80 °C.

### Expression and purification of 6His-hUBA1

Expression of 6His-hUBA1 in Escherichia coli BL21(DE3) cells was performed at large scale due to typically low expression levels. Cultures were supplemented with 1 mM MgSO₄ per liter to support high-density growth and with ampicillin (100 μg/mL). Starter cultures were inoculated at approximately 10 mL per liter of LB medium. Cultures were grown at 37 °C until an OD600 of ∼0.8 was reached (approximately 6–8 h), after which the temperature was reduced to 16 °C. Protein expression was induced with 0.25–0.4 mM IPTG for an additional 16–20 h. Harvested cells were resuspended in 50 mM phosphate buffer, 500 mM NaCl, and 40 mM imidazole, adjusted to pH 7.5. Approximately 24 L of culture yielded ∼300 mL of cell suspension. Cells were passed through a Conti cloth filter and lysed by high-pressure cell disruption (Avestin). MgCl₂ was added to a final concentration of 1 mM together with DNase (0.5 mg/mL). Lysates were clarified by centrifugation (19,000 rpm, 45 min, 4 °C; 45Ti rotor) and filtered through a 0.45 µm syringe filter prior to Ni-NTA chromatography. Clarified lysates were loaded onto a 5 mL HisTrap column pre-equilibrated in the same buffer as the protein at a flow rate of 2.5–4.0 mL/min. After loading, the column was washed with 25 column volumes of the same buffer and hUBA1 was eluted using a 50 mM phosphate buffer, 500 mM NaCl, and 350 mM imidazole, pH 7.5. Eluted hUBA1 was concentrated using a 30 kDa MWCO Amicon centrifugal filter unit and diluted 10x 50 mM Tris and 5 mM β-mercaptoethanol, pH 8.0, to reduce the salt concentration to ∼50 mM. The sample was loaded onto a 5 mL HiTrap Q-HP anion exchange column (Cytiva) and washed with 5 column volumes of Anion Buffer A. Proteins were eluted using a 0–50% gradient of 50 mM Tris, 1 M NaCl, and 5 mM β-mercaptoethanol, pH 8.0, over 40–60 column volumes. Fractions containing hUBA1 were concentrated and further purified by size-exclusion chromatography on a Superdex 200 16/600 column (Cytiva) equilibrated in 25 mM Tris, 300 mM NaCl, and 0.5 mM TCEP, pH 8.0. Homogenous hUBA1 was assessed by SDS-PAGE analysis and Coomassie blue staining. Aliquots were stored at -80°C.

### Expression and purification of untagged ubiquitin

Untagged ubiquitin variants expressed from the pET24a vector were purified using ammonium sulphate precipitation followed by ion exchange and size-exclusion chromatography. Induced bacterial cell pellets were resuspended in a lysis buffer containing 50 mM Tris, 250 mM NaCl, and 0.5 mM TCEP, adjusted to pH 7.5, and supplemented with EDTA-free complete protease inhibitor cocktail (Roche). Cells were lysed by high-pressure cell disruption (Avestin), and insoluble material was removed by centrifugation (27,200 g, 45 min, 4 °C). The clarified lysate was filtered through a 0.2 μm filter, and solid ammonium sulphate was added to 48% saturation for 1 h at room temperature, followed by centrifugation (4,500 rpm, 20 min, room temperature). To precipitate ubiquitin, solid ammonium sulphate was added to the supernatant to 90% saturation and incubated for 1 h at room temperature. The supernatant was discarded, and the pellet resuspended in 2 L of low-salt buffer containing 10 mM ammonium acetate and 0.5 mM TCEP, pH 4.5. Ion exchange chromatography was performed using an ÄKTA Pure system (Cytiva) with a HiTrap SP HP 5 mL column pre-equilibrated in low-salt buffer. After washing with 10 column volumes of low-salt buffer, proteins were eluted using a linear gradient from low-salt to high-salt buffer containing 10 mM ammonium acetate, 0.5 mM TCEP, and 1 M NaCl, pH 4.5. The column was washed with 10 column volumes of high-salt buffer. Fractions containing homogenous ubiquitin were identified by SDS-PAGE, pooled, and concentrated to approximately 10 mg/mL using Vivaspin centrifugal concentrators (Sartorius; 3 kDa MWCO) at 3,000 g and 4 °C. Samples were further purified by size-exclusion chromatography on a HiLoad 26/600 Superdex 75 pg column (GE Healthcare) equilibrated in a buffer containing 50 mM Tris, 150 mM NaCl, and 0.5 mM TCEP, pH 7.5. Fractions containing homogenous ubiquitin were pooled, concentrated to approximately 500 μM, aliquoted, flash-frozen in liquid nitrogen, and stored at −80 °C

### Expression and purification of RNF38 RING and RNF2/BMI1 heterodimer

pGEX4T1 expressing GST-TEV-RNF38 (isoform 1) residues 439-515^22^ was transformed into Escherichia coli BL21 (DE3) Rosetta 2 pLysS (Novagen) chemically competent cells. pGEX4T1 expressing GST-TEV-RNF2 residues 1-114 and pRSFDuet1 expressing 6His-TEV-BMI1 residues 1-109^31^ were co-transformed into the same cells and selected using ampicillin and kanamycin. 12 l of cultures were grown in Luria–Bertani (LB) medium at 37°C to an OD600 of ∼0.8 supplemented with 1 mM MgSO4, 200 μM ZnSO4 was added before induction, and cells were induced with 0.25 mM IPTG at 20°C for 12–16 h. Cell pellets were harvested by centrifugation 3,500 rpm for 30 min at 4 °C. The pellets were resuspended in 25 mM Tris–HCl, 500 mM NaCl, 5 mM DTT, pH 7.5 with 1 mM MgCl2 and DNAseI. Lysis was performed at 12,000 psi using a microfluidizer operating at 4 °C. The lysates were cleared by high-speed centrifugation at 32,000g for 45 min at 4 °C. Then supernatant was filtered through a 0.45-μm syringe filter and loaded on a 20 ml GST-Trap HP column (Cytiva) at 2 ml/min. Once loaded, the UV absorbance reached base line over the 20 cv wash. GST proteins were eluted over 5 cv in 25 mM Tris–HCl, 500 mM NaCl, 2 mM TCEP, 20 mM reduced L-Glutathione, pH 7.8. The elution dialysed overnight in 3.5 MWCO Snakeskin (ThermoFisher) at 4°C in 25 mM HEPES, 350 mM NaCl, 1 mM TCEP, pH 7.5 buffer. To ensure a robust heterodimer, GST-TEV-RNF2/6His-TEV-BMI1 was subjected to a loading and elution buffer (25 mM HEPES, 350 mM NaCl, 350 mM imidazole, 1 mM TCEP, pH 7.5) on a 10 ml His-Trap HP column, then dialyzed with TEV in a separate step. TEV protease and residual GST were removed using a pass back step through GST-Trap and His-Trap columns and the flowthrough was concentrated using 3,500 Da MWCO Amicon centrifugal units. The final purification step was size-exclusion chromatography using a 26/600 Superdex 75 column (Cytiva) pre-equilibrated in 25 mM HEPES, 200 mM NaCl, 1 mM TECP, pH 7.5. Protein purity was assessed using SDS-PAGE and aliquots stored at -80°C.

### Expression and purification of full-length CHIP

pRSF-Duet1 containing human full-length 6His-TEV-CHIP^32^ was transformed into Escherichia coli BL21 (DE3) Rosetta (Novagen) cells and selected with kanamycin. 12 l of cultures were grown in Luria–Bertani (LB) medium at 37°C to an OD600 of ∼0.8 supplemented with 1 mM MgSO4 and cells were induced with 0.25 mM IPTG at 20°C for 12–16 h. Cell pellets were harvested by centrifugation 3,500 rpm for 30 min at 4 °C. The pellets were resuspended in His buffer A (25 mM HEPES, 500 mM NaCl, 2 mM TCEP, pH 7.5) with 1 mM MgCl2 and DNAseI. Lysis was performed at 12,000 psi using a microfluidizer operating at 4 °C. The lysates were cleared by high-speed centrifugation at 32,000g for 45 min at 4 °C. Then supernatant was filtered through a 0.45-μm syringe filter and loaded on a 10 ml His-Trap HP column (Cytiva) at 2 ml/min. Once loaded, the UV absorbance reached base line over the 30 cv wash. 6His-TEV-CHIP was eluted over 6 cv in His buffer B (25 mM HEPES, 500 mM NaCl, 350 mM imidazole, 1 mM TCEP, pH 7.5). The elution was taken for overnight dialysis in 3.5 MWCO Snakeskin (ThermoFisher) at 4°C in 25 mM HEPES, 350 mM NaCl, 1 mM TCEP, pH 7.5 buffer with addition of TEV protease to 1:100 (w/w). TEV protease was removed using a pass back step through a 10 ml His-Trap column and the flowthrough was concentrated using 10,000 Da MWCO Amicon centrifugal unit. This was loaded on a 26/600 Superdex 200 column (Cytiva) pre-equilibrated in 25 mM HEPES, 250 mM NaCl, 1 mM TECP, pH 7.5 buffer. Fractions were checked for purity by SDS-PAGE, concentrated, and aliquots stored at -80°C.

### Preparation of N-maleimido-diazirine–labelled ubiquitin

*N*-maleimido-diazirine^14^ stocks were prepared at 10 mM in anhydrous DMSO and stored in 50 µL aliquots at −20 °C. Ubiquitin cysteine variants were buffer-exchanged into degassed buffer containing 50 mM Tris and 150 mM NaCl, pH 7.0, using Centri Pure Zetadex-25 gel filtration columns (Generon). Labelling reactions were performed at room temperature for 2 h using a five-fold molar excess of N-maleimido-diazirine relative to protein at a protein concentration of 200 µM. Excess reagent was removed by gel filtration using Centri Pure Zetadex-25 columns equilibrated in buffer containing 50 mM Tris, 150 mM NaCl, and 0.5 mM TCEP, pH 7.5. All buffers were degassed, all labelling reactions and labelled proteins were protected from light, and all products were analysed by intact LC–MS.

### NMD-labelled ubiquitin multiple turnover assay

Ubiquitination reactions were carried out by incubating a mixture containing 0.1 μM 6His-UBA1, 0.5 μM UbcH5a, 0.55 μM RNF4, 5.5 μM 4xSUMO-2, 20 μM ubiquitin, 50 mM TRIS, 150 mM NaCl, 5 mM MgCl2, 0.5 mM TCEP, and 0.1% NP-40 at room temperature. The reaction was initiated by adding 3 mM ATP and stopped by addition of reducing SDS-PAGE loading buffer to 2x concentration. Time points were collected at intervals (0, 2, 5, 10, 20, 40, 60, and 100 minutes), the zero-time point taken before ATP addition. Samples were incubated at 95 °C for 5 minutes and analysed using SDS-PAGE on a 10% NuPAGE Bis-Tris polyacrylamide gel with MES buffer, followed by Coomassie Blue staining.

### Preparation of the NMD-Ub-UbcH5a and NMD-Ub-Ubc13 ABPs

UbcH5a (C85K) (25 μM) was incubated with NMD-labelled ubiquitin D32C (100 μM) and 6His–Uba1 (1 μM) at 37°C for 24 h in conjugation buffer (50 mM Tris pH 10.0, 5 mM MgCl2, 1 mM TCEP) ATP (3 mM) was added to initiate the reaction. Homogenous NMD-Ub-E2 was collected using a HiLoad 16/600 Superdex 75 pg column (GE Healthcare) pre-equilibrated with 20 mM HEPES, 150 mM NaCl, 1.0 mM TCEP, pH 7.5. Hamilton syringe (5 mL) was used to inject a 3 mL reaction onto the column. The flow rate was 0.5 mL/ min, fractions were 0.5 mL and homogenous NMD-Ub-E2 was confirmed by SDS-PAGE analysis with Coomassie blue staining. The purified NMD-Ub-E2 was concentrated (Cytiva, Vivaspin, MW cut-off 10 kDa), and stored at -80°C. For the Ubc13-based ABP, Ubc13 C87K, K92A (50 μM) was incubated with NMD-labelled ubiquitin D32C (60 μM) and 6His–Uba1 (0.8 μM) at 37°C for 21 h in conjugation buffer (50 mM Tris pH 10.0, 150 mM NaCl, 5 mM MgCl2, 0.5 mM TCEP). 3 mM ATP was added to initiate the reaction. Homogenous NMD-Ub-E2 was collected using a HiLoad 16/600 Superdex 75 pg column (GE Healthcare) pre-equilibrated with 20 mM HEPES, 150 mM NaCl, 1.0 mM TCEP, pH 7.5. Hamilton syringe (5 mL) was used to inject a 3 mL reaction onto the column. The flow rate was 0.5 mL/ min, fractions were 0.5 mL and homogenous NMD-Ub-E2 was confirmed by SDS-PAGE analysis with Coomassie blue staining. Purified NMD-Ub-E2 was concentrated (Cytiva, Vivaspin, MW cut-off 10 kDa), and stored at -80°C.

### Photocrosslinking of NMD-Ub-UbcH5a or Ubc13 ABPs

Photocrosslinking reactions (40-100 μL) were performed with NMD-Ub-UbcH5a or NMD-Ub-Ubc13/Ube2V2 (∼10 μM) and target protein (10 μM) in reaction buffer (20 mM HEPES, pH 7.5, 150 mM NaCl, 1 mM TCEP). Samples were divided into two portions. One portion was irradiated at 365 nm ∼15 cm away from a 365 nM LED lamp (UHP-T-365-MP, Prizmatix) in an 18-well glass-bottom plate (Ibidi) on an ice-cold metal block for 10 min and the other portion was preserved in the dark on ice. Samples were resolved by SDS-PAGE, 10% NuPAGE Bis-Tris polyacrylamide 1.5 mM gels used to load sample volumes above 20 µL. Proteins were visualized by Coomassie staining.

### MS sample preparation by in-gel tryptic digestion

Proteins were diluted in NuPAGE LDS Sample Buffer (Invitrogen) to 1.2x operating concentration, separated by SDS-PAGE and visualised using filtered Coomassie blue in a sterile dish. The protein bands of interest were cut out of the gel and into ∼1mM cubes. The gel pieces were de-stained overnight in 50mM ammonium bicarbonate (ABC) and 50% Acetonitrile (ACN). Proteins were reduced with 10 mM DTT for 30 mins and alkylated in 50 mM iodoacetamide (VWR) in the dark for 30 mins. Gel pieces were washed with 100mM ABC and then 20 mM ABC, 50% ACN before being dehydrated with 100% ACN for 5 mins. Gel pieces were dried in a fume hood for 15 mins. Trypsin (Thermo) was added (1:100 w:w) according to estimated protein amounts in each band. Trypsin was diluted in 20 mM ABC, 9% ACN and added to the gel pieces in a volume that just covered the pieces (rehydration volume - RV). Gel pieces were incubated for 16 h at 37◦C. To extract the peptides, the rehydration volume (RV) of 100% ACN was added to the gel pieces, followed by RV 5% formic acid, 50% ACN. Pieces were dehydrated again by adding RV 100% ACN. The liquid was evaporated in the gyrovap using the “V-AQ” setting at 45°C. Peptide pellets were resuspended in 0.1% trifluoroacetic acid (TFA) 0.5% acetic acid and submitted for MS analysis. Gel pieces were shaken at RT, 1400 rpm during all steps unless stated otherwise.

### LC-MS/MS peptide analysis

Samples were analyzed on a Thermo Fisher Scientific Lumos Tribrid mass spectrometer coupled with a Thermo Dionex Ultimate 3000 RSLC HPLC. The buffers used for HPLC were 0.1% formic acid as buffer A and 80% acetonitrile (ACN) with 0.08% formic acid as buffer B. Trap column Acclaim pepmap 100 (C5, 100 μM × 2 cm) was used before the main column for sample concentration and cleanup. The peptide samples were loaded onto the trap column using a loading pump with 3% ACN and 0.1% TFA at a flow rate of 5 μl/min. The main column used was EASY- Spray column (C18, 2 μm, 75 μm × 50 cm) with a nano electrospray emitter built in. The flow rate of 300 nl/min was maintained throughout the run. Peptides were separated with a 90 min gradient as detailed within raw data files (see “Data availability” for further details). The separated peptides were analyzed on the mass spectrometer with the following settings. Spray voltage was 2 kV, RF lens level was 30%, and ion transfer tube temperature was 275°C. The mass spectrometer was operated in data-dependent mode with 2 s cycle time. The full scan was performed in the range of 375 to 1500 mass/charge ratio (m/z) at a nominal resolution of 120,000 at 200 m/z and automatic gain control (AGC) was set to 400,000 with a custom maximum injection time of 50 ms. This was followed by the selection of the most intense ions above an intensity threshold of 20000 for higher-energy collision dissociation (HCD) fragmentation, with normalized collision energy set to 30. MS2 scans were acquired for charge states 2 to 7 using an isolation width of 1.6 m/z. MS2 scans were done at resolution 60K using an AGC target of 500,000 and a maximum fill time of 500 ms. Dynamic exclusion was set to 30s.

### Identification of N-maleimido-diazirine-linked peptides

Data analysis was performed with MaxQuant version 2.4.0.0. Default settings were used with a few exceptions. A database of all the recombinant proteins included in the crosslinking assays was used. Digestion was set to Trypsin/P (ignoring lysines and arginine N-terminal to prolines) with a maximum of 8 missed cleavages. Match between runs was enabled. Prior to running the search, two new crosslinkers were added to MaxQuant; Hydrolysed *N*-maleimido-diazirine named ‘NMD169’: Linked composition H(11)O(3)C(8)N, mass 169.0738932246 Da. *N*-maleimido-diazirine named ‘NMD151’: Linked composition H(9)O(2)C(8)N, mass 151.0633285383 Da. Specificity 1 was C, position in peptide 1 was set to anywhere. Protein N-term 1 and C-term 1 were selected. Specificity 2 set for any amino acid (ACDEFGHIKLMNPQRSTVWY), position in peptide 2 was anywhere, protein N-term 2 and protein C-term 2 were selected. To search for crosslinked peptides the crosslinker NMD (non-cleavable) was selected, minimum length for a paired sequence was set to 3 and the maximum peptide mass 12,000 Da. The minimum peptide length for unspecified peptide search was 8 and maximum 25. The search included both intra-protein and inter-protein crosslinked peptides. Oxidation (M), Acetyl (Protein N-term) and Carbamidomethyl (C) were included as variable modifications, with a maximum of 5 per peptide allowed. First search was performed with Oxidation (M) and Acetyl (Protein N-term). Protein and crosslinked peptide level FDR was set to 1%. Filtering of crosslinked peptides is described in Supplementary Datafile 1.

### Intact LC-MS

Intact LC-MS analysis was performed using 0.5 µg of protein injected in buffer containing 50 mM Tris, 150 mM NaCl, and 0.5 mM TCEP. LC-MS measurements were carried out on an Agilent 1200 LC–MS system equipped with a Max-Light Cartridge flow cell and coupled to a 6130 Quadrupole mass spectrometer. An Agilent ZORBAX 300SB-C3 column (5 µm, 2.1 × 150 mm) was used unless otherwise stated. Protein elution was monitored by UV absorbance at 214 and 280 nm. Mass spectra were acquired in positive ion mode, and intact protein masses were determined by deconvolution using MS ChemStation software (Agilent Technologies).

## Supporting information

Supplementary Datafile 1

## Acknowledgements

We acknowledge the technical and research staff of the Ciulli Laboratories for the set-up and upkeep of protein expression and purification infrastructure at CeTPD. Many thanks to Robert Gourlay and colleagues from the University of Dundee MRC proteomics facility for MS data acquisition. Thanks to Satpal Virdee (University of Dundee) for assistance with in tact LC-MS analysis.

## SUPPLEMENTARY FIGURE LEGENDS

**Supplementary Figure 1.**
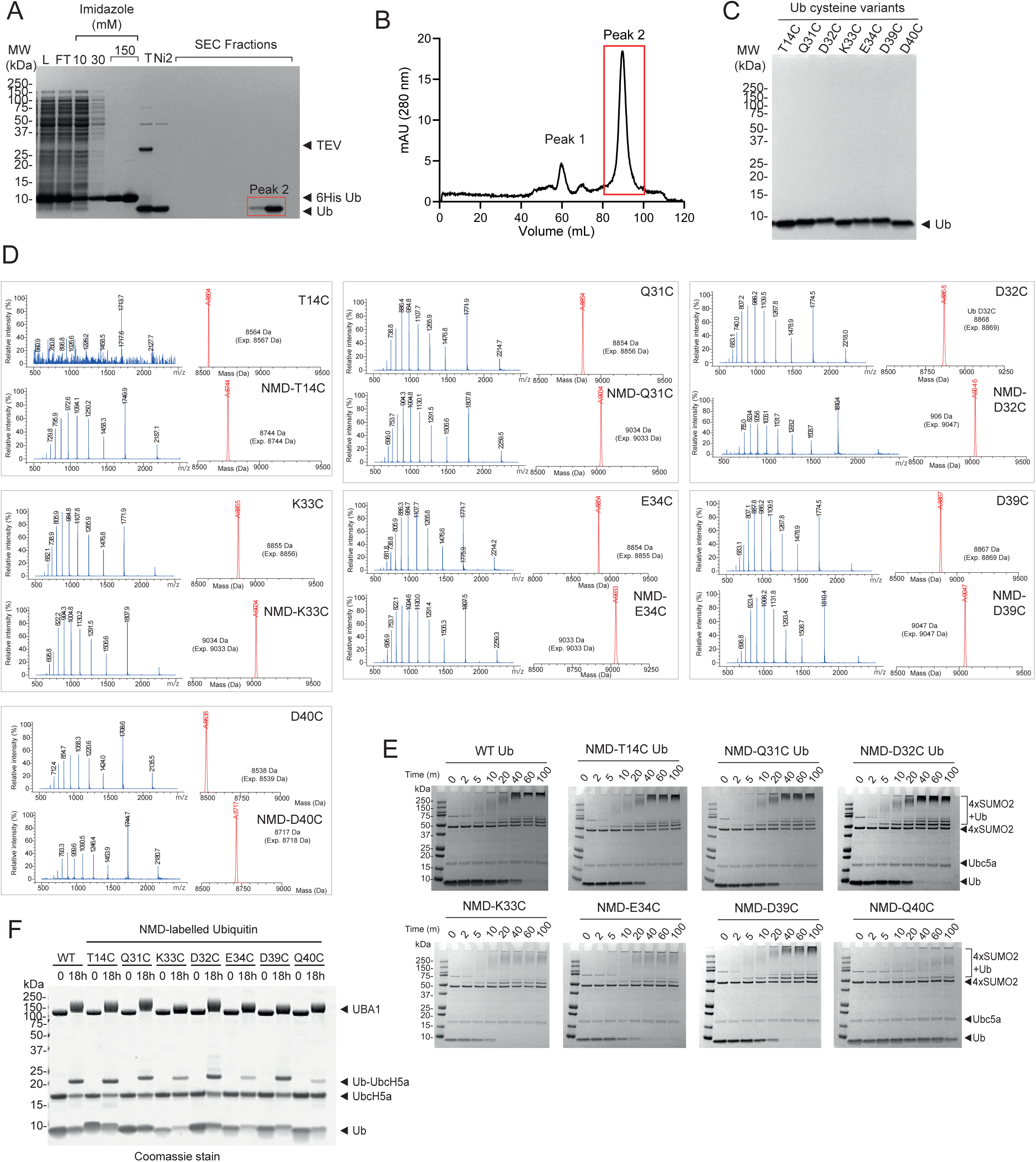
Purification, NMD labelling and conjugation to UbcH5a of seven ubiquitin mutants. Related to Figure 1. A. Coomassie stained SDS-PAGE gel of protein samples taken during recombinant ubiquitin protein expression and purification (L = Nickel-NTA column load, FT – Nickel-NTA column flow through, 10 – 10mM imidazole wash, 30 – 30mM Imidazole wash, 150 – 150mM imidazole elutions, T – Purified 6His-Ub + TEV, Ni2 – Post TEV Nickel-NTA column flow-through, SEC – size exclusion chromatography. B. 280nm absorbance trace of SEC elution. C. Coomassie stained SDS-PAGE gel of 2 µg each purified cysteine ubiquitin variant. D. LC-MS for unlabelled and NMD-labelled ubiquitin variants as indicated. E. Conjugation of NMD-Ub variants (20 µM) to 4xSUMO-2 (5.5 µM) in the presence of RNF4 (0.55 µM), UbcH5a (0.5 µM), and UBA1 (0.1 µM). Samples taken after the indicated time points post ATP addition were analysed by Coomassie-stained SDS-PAGE. F. Conjugation of WT ubiquitin and NMD-Ub variants by UBA1 to UbcH5a C85K to form the NMD-Ub-UbcH5a ABPs.

**Supplementary Figure 2.**
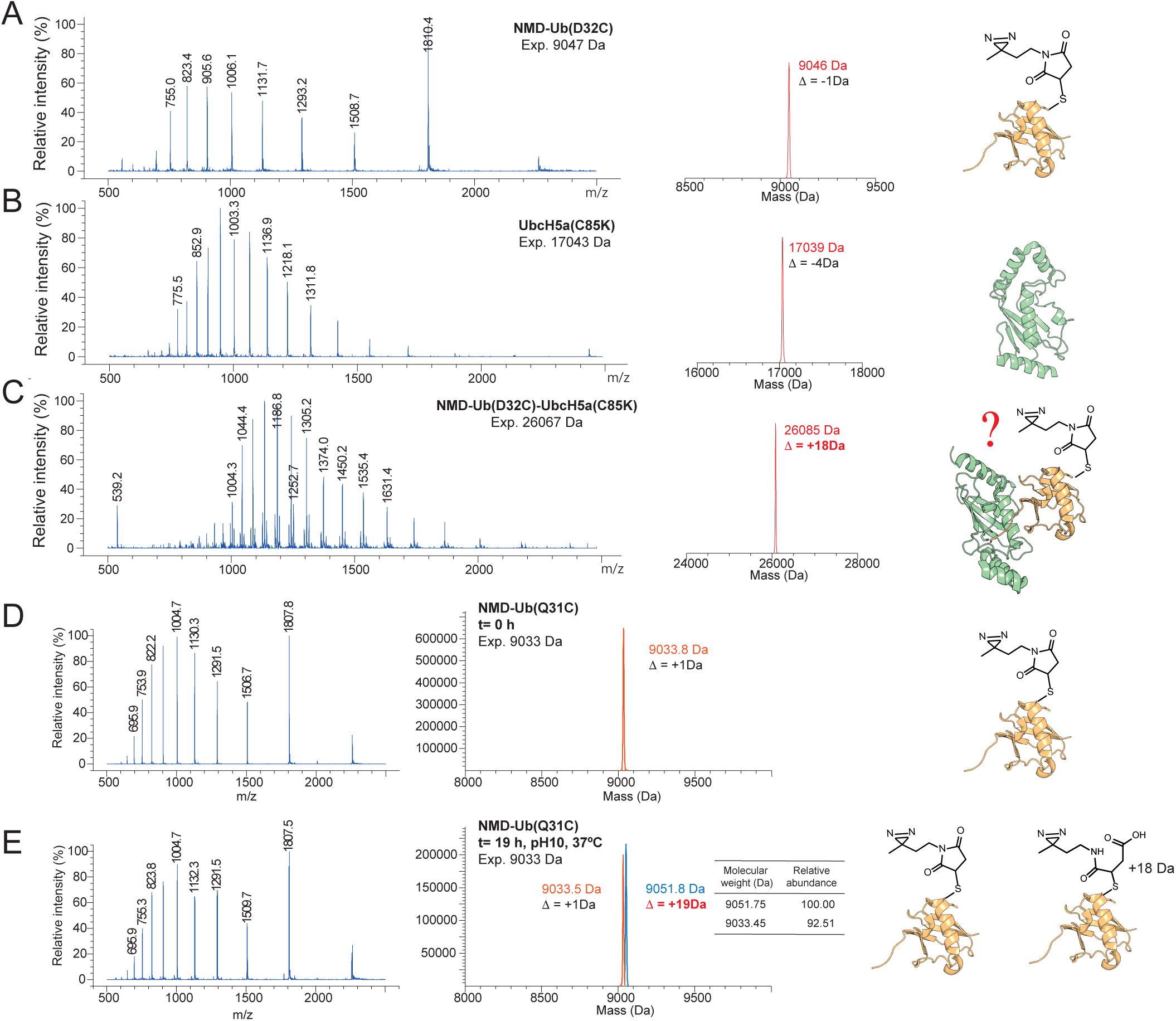
Hydrolysis of NMD under conjugation assay conditions results in an 18 Da increase in mass of NMD-Ubiquitin. Related to Figure 2. A-C. LC-MS analysis of NMD-labelled ubiquitin (D32C) (A), UbcH5a (C85K) (B) and NMD-Ub-UbcH5a (C). The expected mass for NMD-Ub-UbcH5a accounts for the loss of 18 Da during the formation of the isopeptide bond between Ub G76 and UbcH5a C85K. D+E. LC-MS analysis of NMD-Ub(Q31C) before (D) and after (E) incubation under the conditions of the ubiquitin conjugation assay.

**Supplementary Figure 3.**
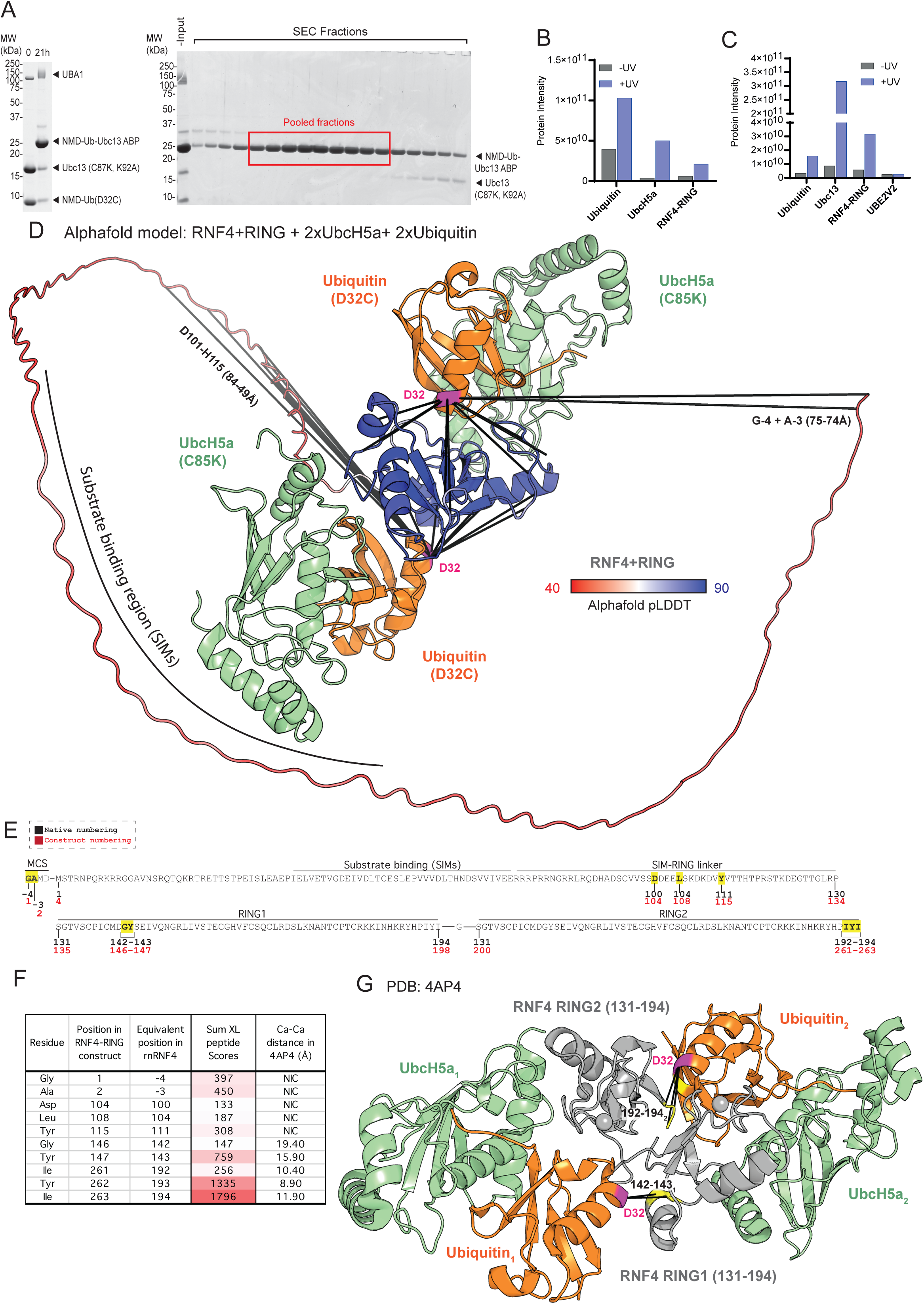
UV-dependent high molecular weight adducts are indicative of cross-linking of NMD-Ub-Ubch5a and NMD-Ub-Ubc13 to RNF4+RING. Related to Figure 2. A. Conjugation of NMD-Ub to Ubc13 with samples taken prior to ATP addition (“0”) or after 21 hours incubation at 37°C were analysed by SDS-PAGE (left). NMD-Ub-Ubc13 was purified by superdex 75 SEC (right) and pooled fractions indicated. B+C. Total peptide (protein) intensity values (sum of all non-cross-linked peptide intensities) for the indicated proteins in UV-specific species excised from the gels shown in Figure 2C and Figure 2D respectively. D. Alphafold model generated from hRNF4(1-190) and RNF4+RING(127-190) with two copies each of UbcH5a-C85K) (pale green) and ubiquitin (D32C) (orange). The RNF4 construct is coloured by pLDDT and long-distance cross-links annotated. E. Sites of crosslinking to the RNF4+RING construct using the NMD-Ub-UbcH5a ABP. Distances according to the UbcH5a-RNF4+RING structure (PDB: 4AP4) are summarised in F and shown in G. NIC – Not in the construct used for the structural study. NM – Present in the construct used in the study but not modelled in the final structure.

**Supplementary Figure 4.**
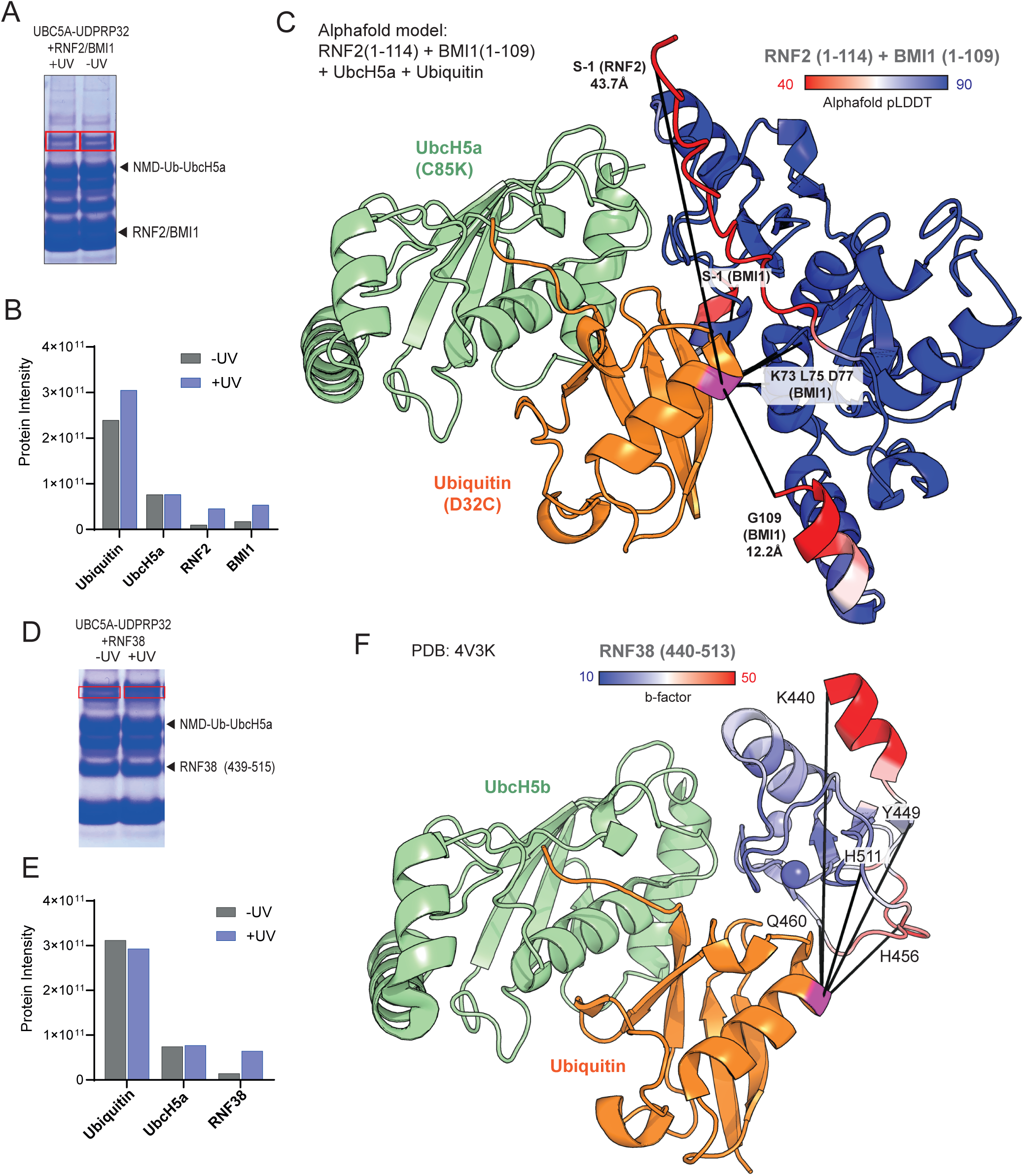
Crosslinking of NMD-Ub-UbcH5a to RNF2/BMI1 and RNF38. Related to Figure 3. A. SDS-PAGE analysis of UV irradiated (+UV) or non-irradiated (-UV) reactions containing RNF2(1-114)/BMI1(1-109) with NMD-Ub-UbcH5a. Identified regions (red box) were excised for digestion and LC-MS/MS analysis. B. Total peptide (protein) intensity for the indicated proteins determined from non-crosslinked peptides determined by LC-MS/MS analysis of samples from A. C. Alphafold model of UbcH5a(C85K), Ubiquitin(D32C) (orange) and RNF2(1-114)/BMI1(1-109) coloured by pLDDT scores. Identified cross-links are indicated. D. SDS-PAGE analysis of UV irradiated (+UV) or non-irradiated (-UV) reactions containing RNF38(439-515) and NMD-Ub-UbcH5a. Identified regions (red box) were excised for analysis by LC-MS/MS. E. Total peptide (protein) intensity for the indicated proteins as determined by LC-MS/MS. F. Cross-links from ubiquitin residue 32 to RNF38(440-513) in the structure PDB 4V3K. b-factor values are colour coded as in the key.

**Supplementary Figure 5.**
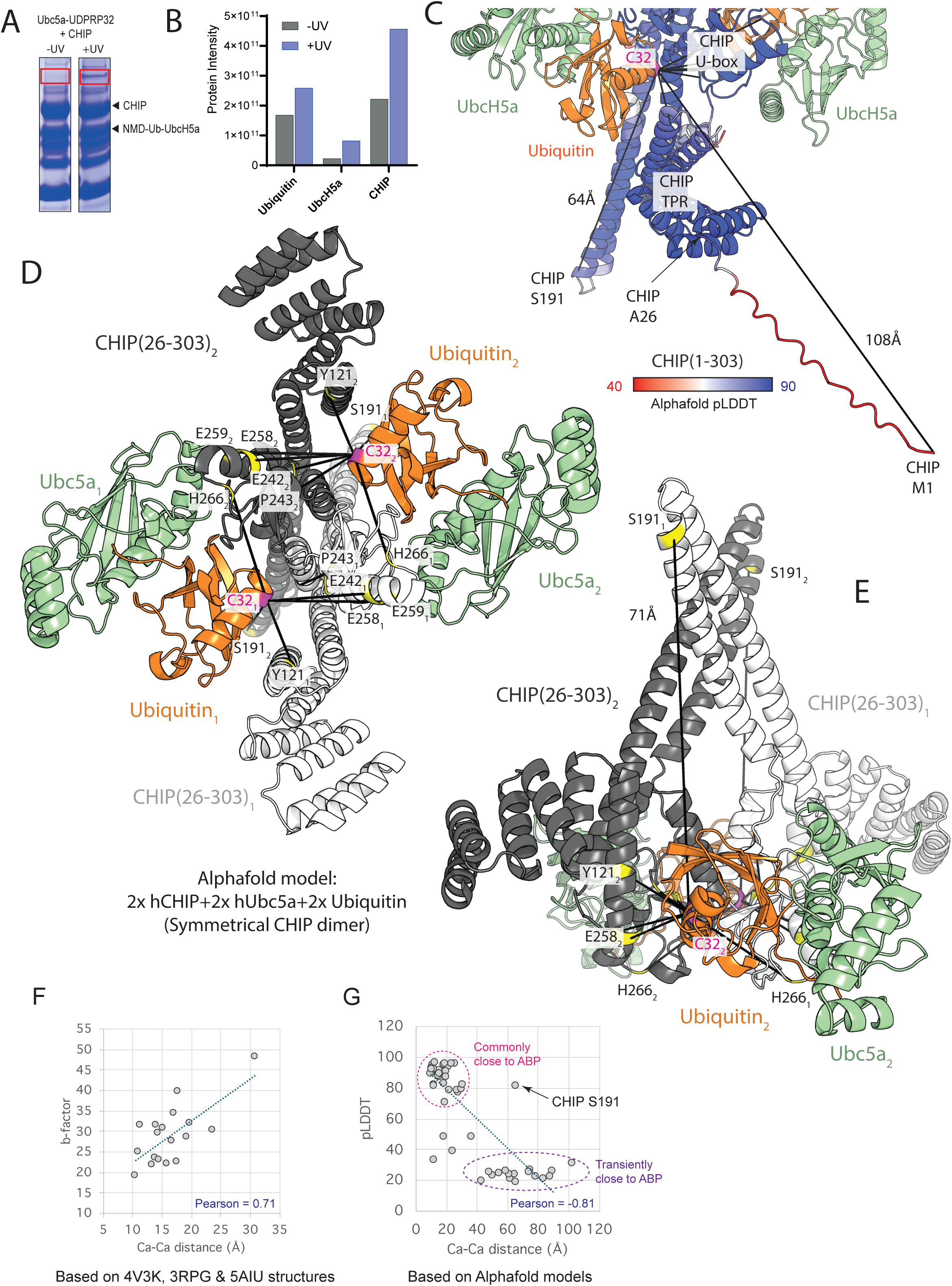
A symmetrical dimer model for CHIP binding Ub-E2. Related to figure 4. A. SDS-PAGE analysis of UV irradiated (+UV) or non-irradiated (-UV) reactions containing full-length human CHIP with NMD-Ub-UbcH5a. Identified regions (red box)were excised from both lanes for digestion and LC-MS/MS analysis. B. Total peptide (protein) intensity for the indicated proteins determined from non-crosslinked peptides determined by LC-MS/MS analysis of samples from A. C. Alphafold model of UbcH5a(C85K) (pale green), Ubiquitin(D32C) (orange) and two copies of full length human CHIP (coloured by pLDDT scores). Identified cross-links are indicated. D, E. Alphafold model for a 2xUbcH5a, 2xubiquitin, 2xhCHIP(26-303) symmetrical dimer. Identified crosslinks are indicated. D is viewed along the axis of symmetry with the helical bundles projecting into the page, and E shows a side-view with one Ubiquitin-UbcH5a complex in the foreground. F. Relationship between measured crosslinker Cα-Cα distance and b-factor of the E3 ligase residue according to the structures shown in Figures 2J, 3D and 3H/S4F. G. Relationship between measured crosslinker Cα-Cα distance and Alphafold pLDDT values of the E3 ligase residues according to the models shown in Figures 4E, S3D and S4C. Pearson correlation coefficients are indicated in G and H.

